# Lung infection by *P. aeruginosa* induces neuroinflammation and blood-brain barrier dysfunction in mice

**DOI:** 10.1101/2023.01.23.524949

**Authors:** Nuria Villalba, Yonggang Ma, Sarah A. Gahan, Aurelie Joly-Amado, Sam Spence, Xiaoyuan Yang, Kevin Nash, Sarah Y. Yuan

## Abstract

**Background:** Severe lung infection can lead to brain dysfunction and neurobehavioral disorders. The mechanisms that regulate the lung-brain axis of inflammatory response to respiratory infection are incompletely understood. This study examined the effects of lung infection causing systemic and neuroinflammation as a potential mechanism contributing to blood-brain barrier (BBB) leakage and behavioral impairment.

**Methods:** Pneumonia was induced in adult C57BL/6 mice by intratracheal inoculation of *Pseudomonas aeruginosa* (PA). Solute extravasation, histology, immunofluorescence, RT-PCR, multiphoton imaging and neurological testing were performed in this study.

**Results:** Lung infection caused alveolar-capillary barrier injury as indicated by leakage of plasma proteins across pulmonary microvessels and histopathological characteristics of pulmonary edema (alveolar wall thickening, microvessel congestion, and neutrophil infiltration). PA also caused significant BBB dysfunction characterized by leakage of different sized molecules across cerebral microvessels and a decreased expression of cell-cell junctions (VE-cadherin, claudin-5) in the brain. BBB leakage peaked at 24 hours and lasted for 7 days post-inoculation. Additionally, mice with lung infection displayed hyperlocomotion and anxiety-like behaviors. To test whether cerebral dysfunction was caused by PA directly or indirectly, we measured bacterial load in multiple organs. While PA loads were detected in the lungs up to 7 days post-inoculation, bacteria were not detected in the brain as evidenced by negative cerebral spinal fluid (CSF) cultures and lack of distribution in different brain regions or isolated cerebral microvessels. However, mice with PA lung infection demonstrated increased mRNA expression in the brain of pro-inflammatory cytokines (IL-1β, IL-6, and TNF-α), chemokines (CXCL-1, CXCL-2) and adhesion molecules (VCAM-1 and ICAM-1) along with CD11b+ cell recruitment, corresponding to their increased blood levels of white cells (polymorphonuclear cells) and cytokines. To confirm the direct effect of cytokines on endothelial permeability, we measured cell-cell adhesive barrier resistance and junction morphology in mouse brain microvascular endothelial cell monolayers, where administration of IL-1β induced a significant reduction of barrier function coupled with tight junction (TJ) diffusion and disorganization. Combined treatment with IL-1β and TNFα augmented the barrier injury.

**Conclusions:** These results suggest that lung bacterial infection causes cerebral microvascular leakage and neuroinflammation via a mechanism involving cytokine-induced BBB injury.

## Introduction

Severe lung injury caused by bacterial infections may contribute to clinical complications such as multiple organ dysfunction, including cerebral and neurological disorders, which have been associated with high morbidity and mortality (1–3). Several factors, from pathogen load to duration of infection, determine patients’ outcome with neuroinflammation as the link between long-term neurological damage and brain dysfunction resulting in a myriad of cognitive complications and behavioral symptoms.

The importance of lung-brain communication has been recognized based on strong clinical and experimental evidence. For example, clinical studies have demonstrated that patients with pneumonia often experience changes in mental status, such as confusion, low alertness and even delirium, specifically in older patients (4). Additionally, community-acquired pneumonia, the most prevalent cause of sepsis, can also induce acute and chronic changes in the central nervous system (CNS) (5–8). Sepsis-associated encephalopathy (SAE), a well-characterized state of long-term cognitive impairment and neurological dysfunction develops in approximately 70% of sepsis survivors (9). The causes of SAE are not completely understood albeit one of the explanations could be that a compromised BBB leads to brain dysfunction and CNS pathology (10, 11). A broad range of pathogens are capable of permanently or temporary affect brain functions (12, 13). Unlike viruses, which can infiltrate the CNS, direct dissemination of bacteria into the brain is rare and, therefore, other factors must contribute to the cerebral pathophysiology.

In the present study, we used a clinically relevant mouse model of bacterial pneumonia. We inoculated freshly prepared PA intratracheally into mice and examined the relationship between lung infection and cerebral dysfunction. Specifically, we tested whether the changes in the brain are due to a direct effect of bacteria penetrated the CNS, or indirectly via systemic and neuroinflammation triggered by pulmonary infection.

## Methods

### Bacteria growth and inoculum preparation

PA was selected for this study because it is known to be the most common cause of nosocomial pneumonia (14). PA (PAO1 strain) (ATTCC; #27853) was cultured following manufacturer’s instructions. Prior to infection, freshly cultured bacteria from frozen aliquots (−80 °C) were grown overnight at 37 °C in tryptic soy broth (Remel; #R455052) with vigorous agitation (180 rpm) and harvested in log phase to an optical density at 600-nm (OD_600_) of 0.3 (only for survival studies, mice were inoculated with OD_600_ = 0.6). Bacteria were centrifuged at 3,000 rpm for 10 min, rinsed and resuspended in sterile phosphate-buffered saline (PBS, pH 7.4). Serial dilutions and colony count of the inoculum were performed to obtain the number of colony-forming units (CFUs) per mL (CFU/mL). For histology experiments and multiphoton imaging, PA labelled with a green fluorescent protein gene (PA-GFP; ATCC; #10145GFP) was used. PA-GFP was cultured in nutrient agar (BD Difco^™^; #213000) or broth (BD Difco^™^; #234000) supplemented with 300 μg ampicillin following manufacturer’s instructions as described above and then inoculated into mice.

### Animal model of pneumonia

Male and female C57BL/6 mice between 8 and 16 weeks of age were used in all experiments. All mice were housed in a temperature and humidity-controlled housing with a 12-hour light:dark diurnal cycle. Littermates of the same age were allocated to experimental groups. All experimental animal protocols were approved by the University of South Florida Institutional Animal Care and Use Committee (IACUC; protocol 8421R) and were conducted in accordance with the Guide for Care and Use of Laboratory Animals.

Under anesthesia, forty-microliters of bacteria reconstituted in sterile PBS [OD_600_ = 0.3; 2×10^7^(CFUs/mL)] were directly injected into the mouse lungs by intratracheal instillation using a 22-G cannula via the oropharynx using a mouse intubation kit (Hallowell EMC, MA) and an otoscope. Control group received the same volume of sterile PBS (uninfected group). After inoculation, mice were kept in a supine position for a few breaths or at least 30 seconds. Once awake, mice were returned to their cages, monitored regularly and allowed access to food and water. For clinical relevance, antibiotics (imipenem+cilastatin formulated in 0.9% saline solution) were administrated subcutaneously at a dose of 5 mg/Kg starting at 6 hours post-infection. The general conditions of each mouse were monitored over a period of 7 days. All procedures were performed in a bio-safety cabinet in a designated bio-safety level-II room according to the Biosafety protocols approved by the University of South Florida.

### Measurement of microvascular permeability *in vivo*

The leakage of circulating solutes across microvessels was evaluated by measuring tissue (brain and lung) accumulation of tracer molecules of different sizes using the ratiometric assay and near-infrared imaging (NIR) as we previously described (15). Time-course of vascular permeability changes was assessed in anesthetized mice intravenously injected through the retro-orbital venous sinus with sodium fluorescein (NaFl; 5 μL/g of a 100 mg/mL NaFl solution; 376 Da; Sigma-Aldrich; #6377) at 24 hours, 7 days and 1-month post-infection. Then, mice were anesthetized with urethane (1.75 mg/Kg; intraperitoneally) and transcardially perfused with ice-cold 0.1 M PBS. After dissection, brains and lung lobes were harvested, weighed and homogenized. Tissue homogenates were centrifuged at 12,000 × g for 20 min at 4 °C. The fluorescence of a 100 μL aliquot of the brain and lung samples along with 100 μL of blank or standards was measured on a fluorescence microplate reader (SpectraMax M3; Molecular Devices, Sunnyvale, CA). The concentrations of the samples were within the linear range of the standard curve. Amount of dye (ng) was normalized per pg or μg protein in the extract.

In a different set of experiments, permeability to Alexa Fluor 70-kDa (5 μg/g of a 1 mg/mL solution; Sigma-Aldrich; #R9379) was assessed in the lungs of controls and infected animals at 24 hours post-infection. For NIR of lungs (left lobe), animals received albumin (Albumin-CF^®^680 dye; ~ 70-kDa; 5 μg/g of a 1 mg/mL solution; Biotum; #20292). Tracer was allowed to distribute in freely moving mice for 4 hours and then animals were cardially perfused with PBS. For NIR imaging of brains, anesthetized animals received a combined injection of 10-kDa dextran (10 μg/g of a 1 mg/mL solution; CF^TM^790-10-kDa dextran; #80121; Stellar Scientific) and albumin (Albumin-CF^®^680 dye) intravenously. Organs (left lung lobe and whole brain) were imaged at 700 and 800 nm with a near-infrared imaging system (Odyssey CLx; LI-COR Biosciences, Lincoln, NE).

### Measurements of bacteria load in tissues and CSF

Mouse lungs and spleen were harvested at different time points after infection, weighted and then homogenized in tryptic soy broth under sterile conditions. One hundred μL of the homogenate were plated on tryptic soy agar plates for quantification of colonies. CSF samples were obtained under anesthesia through the cisterna magna using a custom-made glass cannula. Ten-fold serial dilutions of the homogenates and CSF were prepared and spread onto agar plates. Next day, CFUs were counted. For the identification of bacteremia (qualitatively), 100 μL of blood were obtained under sterile conditions and cultured in culture vials (BD Bactec^TM^ peds plus culture vials; #442020) for 4 days. Then, 100 μL of the culture vials were streaked onto a 5% sheep blood in tryptic soy agar plates (Hardy diagnostics; #A10) for colony counts.

### Proinflammatory mediators in plasma and brain tissue

Plasma samples obtained from blood collected by cardiac puncture in heparin tubes (BD Microtainer^®^ blood collection tubes; BD Biosciences; #365965) were analyzed for IL-6, IL-1β and TNF-α using indirect sandwich enzyme-linked immunosorbent assay (ELISA) (Sigma-Aldrich; #RAB0308 and #RAB0477) according to the manufacturer’s instructions.

Total RNA was extracted from brain cortices and hippocampi from PA-infected and control animals using TRIzol^®^ Reagent (Invitrogen; #15596) and RNeasy Plus kit (Qiagen). RNA concentration was measured using NanoDrop One Spectrophotometer (Thermo scientific). Reverse transcription (RT)-PCR of RNA was performed using High-Capacity RNA-to-cDNA Kit (Thermo Fisher Scientific; #4387406). cDNA was amplified for real-time detection with TaqMan^™^ Gene Expression Master Mix (Thermo Fisher Scientific; #4369016) plus individual primers for IL-1β, IL-6, TNF-α, ICAM-1, VCAM-1, CXCL1 and CXCL2 (FAM tagged; Thermo Fisher Scientific) on a Biorad CFX Connect^™^ machine. Expression data were normalized to Gapdh mRNA levels. The data were calculated as 2(Ct(Gapdh–gene of interest)) to compare infected group to controls. MIQE guidelines were followed for all the PCR experiments and analysis (16).

### Flow Cytometry

Mice were euthanized and perfused with sterile PBS through the left ventricle of the heart. The brains (400-500 mg; without the cerebellum) were harvested, cut along the coronal plane with a razor into three pieces and digested into cell suspensions using a Multi Tissue Dissociation kit 1 (Miltenyi Biotec, Germany; #130-110-201) and a gentleMACS^™^ OctoDissociator with heaters (Miltenyi Biotec; #130-096-427) following manufacturer’s instructions. During the final steps, myelin and erythrocytes were removed using a neural debris removal kit (Miltenyi Biotec; 130-109-398) and red cell lysis solution (Biolegend; #420301), respectively. Total cell count in each brain was quantified using a Luna-II automated cell counter. Cells were then subjected to antibody staining for flow cytometry analysis. The cell suspension was stained with a panel of antibodies against CD45-Alexa 700 (1:1200; Biolegend; #147716) and CD11b-PE dazzle (1:800; Biolegend; #101256). A corresponding isotype IgG antibody was used as negative staining control. Dead cells were excluded using the live/dead stain Zombie NIR fixable viability dye (1:2000; Biolegend; #423105). The samples were run on a Cytek^®^ Northern Lights^™^ cytometer (Cytek biosciences) and data were analyzed using FlowJo (FlowJo LLC, Ashland, OR). Cell populations were calculated based on relative abundance obtained by flow cytometry (number = total cell count x % of positive cells) (17). Cells were gated using FSCA and FSCH to identify singlets, then gated for live cells by taking a population that was negative for live/dead stain. Antibodies concentrations were chosen based on previous antibody titrations to determine the dilution that provides the best separation of cell populations. Each “n” refers to a single brain obtained from a single mouse, so that each n represents a different mouse. Brains were not pooled together for these experiments. See Suppl. Fig. 1 for gating strategy used.

### Endothelial cell culture

Primary mouse brain microvascular endothelial cells (Cell Biologics, Chicago, IL; #C57-6023) were grown in the recommended culture medium (Cell Biologics; #M1168) supplemented with 10% (v/v) fetal calf serum at 37 °C and 5% CO2, split at a ratio of 1:2 and used at passage two. Endothelial cell monolayers were used for barrier integrity and immunofluorescence experiments.

### Assessment of barrier function by electric cell-substrate impedance sensing (ECIS) assay

Barrier integrity of brain microvascular endothelial cells was performed using a protocol previously described by our group (15, 18). Cell monolayers were grown to reach confluence in order to obtain a polarized endothelium as evidenced by high and stable transendothelial electrical resistance (TER, ohm) values (15). Mouse brain endothelial cell barrier function was determined by measuring the cell-cell adhesion barrier resistance to electric current using an ECIS system (Model Zθ Applied BioPhysics Inc., Troy, NY). Briefly, 250 μL of cell suspension (5 Å~ 105 cells per mL) were seeded to each well of two ECIS culture arrays consisting of 8-wells with 10-gold microelectrodes (8W1E+) following manufacturer’s instructions. Resistance was recorded in real time with the following settings: alternating current (1-volt) and 4-kHz frequency at 7-second intervals. Data were normalized to baseline measurements just prior to the onset of treatment (t=0) and expressed as resistance change (ohms). Both data acquisition and processing were performed using the ECIS Z-theta Analysis Software supplied by Applied Biophysics to determine the resistance (ohm @ 4kHz) values reflecting cell monolayer integrity and overall response to cytokines. In a different set of experiments, cells were exposed to IL-1β (Miltenyi Biotec; #130-094-053) and IL-1β + TNF-α (Biolegend; #575204). Cytokines were dissolved in PBS + 0.1% BSA at 2, 20 and 200 ng/mL. Vehicle control was diluent alone at the same volume as treatment. Data were normalized to baseline, and comparisons were made between cytokine treatment and vehicle control. Data show mean ± S.E.M. (n=3).

### Immunocytochemistry

Brain endothelial cell monolayers were plated on Lab-Tek^®^ chamber slides (Thermofisher Scientific; #1777399PK) and exposed to cytokines (IL-1β, 20 ng/mL), washed in PBS containing 2 mM CaCl_2_ and 2 mM MgCl_2_ and then fixed in 2% paraformaldehyde (PFA) in PBS for 5 min followed by 4% PFA for 15 min. Cells were washed in PBS twice, permeabilized for 10 min (in 0.1% Triton-X/PBS) and blocked (10% normal donkey serum in 3% BSA/PBS) for 1 h at room temperature. Slides were incubated with ZO-1 primary antibody (1:100; Thermofisher; #40-2200) overnight at 4 °C. Next day, slide chambers were washed seven times and incubated with appropriate secondary Alexa Fluor^®^ antibodies (diluted at 1:500). Chambers were carefully removed, and slides were then cover slipped using ProLong^™^ diamond antifade mountant with DAPI (Life Technologies; #P36966).

### Tissue processing and histology

Tissues (brain and lung) harvested from animals 24 hours after infection with GFP-labelled PA were subjected to multiphoton imaging. Mice were anesthetized and injected intravenously with 200 μL of Evans blue solution containing 3% Evans blue fluorescent dye (Sigma-Aldrich; #E2129) in 5% BSA/PBS to visualize vasculature prior to organ extraction and ex-vivo multiphoton imaging.

For visualizing PA distribution in tissue by confocal microscopy, mice infected with GFP-labelled PA and controls were anesthetized and intracardially perfused through the left ventricle with PBS and 4% PFA thereafter. Organs (lungs, spleen and brains) were post-fixed in 4% PFA overnight, rinsed and preserved in 10% sucrose solution in PBS for one day or until the tissue sank to the bottom. Mouse tissues were collected and embedded in Tissue Plus^®^ O.C.T. compound (Fisher HealthCare; #4585) in disposable base molds (Fisher HealthCare; #22363553). Blocks were then frozen in liquid nitrogen and stored in −80 °C until sectioned in a cryostat. Frozen blocks were cut into sections (15 μm thick) on a Leica CM1950 cryostat and slides were stored at −20 °C until use. On the day of the experiment, slides were thawed in the hood overnight and stained with Dylight 488-conjugated Lycopersicum esculentum lectin (1:100; Vector laboratories; #DL-1174) to visualize brain microvessels. Slides were mounted in ProLongTM diamond antifade mountant with DAPI.

Brain microvessels from control and PA-infected mice were freshly isolated from brain cortices as previously described (19). Briefly, pial vessels were carefully removed, and the remaining tissue was homogenized with a loose-fitting Dounce homogenizer in MCDB 131 medium. Homogenates were then centrifuged for 5 min at 2,000g at 4 °C. Pellets were resuspended in 15% (wt/vol) dextran-PBS and centrifuged again at 10,000g for 15 min at 4 °C. Myelin was carefully removed from the tube and the pellet was removed with 1 mL DPBS and passed through a 40 μm strainer. The strainer was reversed and washed with 10 mL MCDB 131 media to retrieve the microvessels. Microvessels were fixed in 2% PFA and cytospinned onto microscope slides.

For immunohistochemistry, slides were permeabilized and blocked for 1 hour at RT in 10% normal donkey serum diluted with PBS/3% BSA and 0.3 % Triton-X. Slides were incubated overnight with the primary antibodies against neutrophils (1:100; Cederlane; #CL8993AP). Lectin was used to label blood vessels. All antibody dilutions used were optimized by titration.

For visualizing lung structure, hematoxylin and eosin (H&E) staining was performed. Animals were transcardially perfused via the left ventricle with PBS first and then 4% PFA. After one day of post-fixation in the same fixative, lungs were embedded in paraffin using an automatic tissue processor (Thermo Scientific) and then sectioned (4 μm) using a microtome. Slides were deparaffinated and then stained with H&E following standard procedures. All slides were imaged at the same time and using the same settings with a brightfield Olympus VS120 slide scanning imaging system (x20 objective).

### Microscopy and Imaging

Confocal microscopy was performed using two different lenses ×20 HC PL APO (0.75NA [numerical aperture], WD [working distance] 0.62 mm) and a ×63 HC PL APO (Oil immersion, 1.4NA, WD 0.14 mm). Images were obtained as z-stacks using Leica STED SP8 Laser Confocal Microscope and image software (LAS X). For confocal microscopy, GFP-labelled PA was excited with the 488-nm laser.

Two-photon imaging was performed using an Olympus FV1000 MPE microscope equipped with 4 spectral detectors, a x25 XL Plan N 1.05 N.A. water immersion objective and a Coherent Chameleon Ultra II IR laser. For multiphoton imaging, GFP-labelled PA was excited with 924-nm laser pulses, emission was detected at 500–520 nm. Evans blue was excited at 945-nm and detected at 620–750-nm. The raw 3D images were reconstructed from z-stacks using Imaris software 9.7 (RRID:SCR_007370).

### Open field behavioral test

Spontaneous locomotor activity and anxiety-like behavior were measured by open field test assessed at 1-week after PA infection (20). Mice were placed in an open field chamber and were allowed free exploration for 15 min. The distance traveled was recorded and analyzed using ANY-maze^®^ video tracking system (Stoelting Co, IL, USA). Increased overall distance travelled in the chamber and time spent in the center were considered an index of greater locomotor activity and anxiety-like behavior.

### Hematology and biochemistry

White blood cell counts, cell differentials (neutrophils and monocytes) and total protein were measured in mouse blood collected in heparin tubes using a ProCyte Dx hematology analyzer (IDEXX, Westbrook, Maine). Blood pH was measured in fresh blood using an iSTAT device (Abbott, Hightstown, New Jersey).

### Statistical analysis

Data were analyzed using GraphPad Prism (version 9.4.0; GraphPad Software Inc., San Diego, CA). Reported values are expressed as mean ± standard error of the mean (SEM). For animal experiments, female and male mice were randomly selected and used as biological replicates. Normality of the data was tested using the Shapiro–Wilk test. For data that were not normally distributed (nonparametric data), the Mann–Whitney test (two groups, one variable) or Kruskal–Wallis test followed by Dunn’s correction (>2 variables) was used. Data with normal distribution were analyzed by a two-tailed unpaired Student’s t–test (one variable) or one–way ANOVA with Dunnett’s or Tukey’s correction (>2 variables). Statistical test used to analyze the data is described in the figure legends. In all tests a 95% confidence interval was used. Figures show exact P values.

## Results

### Effect of PA inoculation on mouse mortality and lung injury

PA-induced pneumonia was associated with mouse mortality, which depended on the inoculum’s load or number of infecting bacteria. Survival studies were performed using two inoculum’s loads, OD_600_ of 0.3 (~2×10^7^ CFUs/mL) and OD_600_ of 0.6. Infected mice with PA at an OD_600_ of 0.3 and OD_600_ of 0.6 (~9×10^8^ CFUs/mL) induced a 16% and 50% mortality at day 7, respectively, compared to 100% survival in the control group (Suppl. Fig 2a). All infected mice (OD_600_ 0.3) showed lethargy, hypothermia and remained immobile or moved slowly in the cage starting at 4 hours post-infection (Suppl. Fig. 2b). Those who did not die within 24 hours showed significant signs of improvement in mobility and appearance. Additionally, PA also caused acidosis and high levels of proteins in the blood (hyperproteinemia), both metabolic parameters associated with active infection (Suppl. Fig 2c,d). Together, these data support the validity of the PA-induced pneumonia model.

Next, we assessed the pulmonary vascular barrier in vivo in PA infected mice (OD_600_ 0.3; 24 hours post-infection) and uninfected controls by measuring extravasation of intravenously injected 70-kDa fluorescent tracer. Upon PA challenge, plasma leakage from lung microvessels, indicated by higher interstitial accumulation of 70-kDa dextran, was detected in infected mice compared with that in controls (Fig. 1a,b). PA infection also caused a significant protein leakage (Fig. 1c). Lung transvascular total protein flux increased by approximately 8-fold, peaked at 6 hours post-infection and started to decrease by day 7 post-infection. H&E staining of lung sections post-infection showed that PA evoked a robust inflammatory response (Fig. 1d). Compared to control lungs characterized by thin alveolar walls and clear alveolar airspaces with few number of neutrophils confined to the septal network, lung sections obtained from animals inoculated with PA were characterized by congested vasculature, thickened alveolar walls with the presence of microthrombi and alveolar infiltration of neutrophils (Fig. 1d,e).

**Figure 1.**
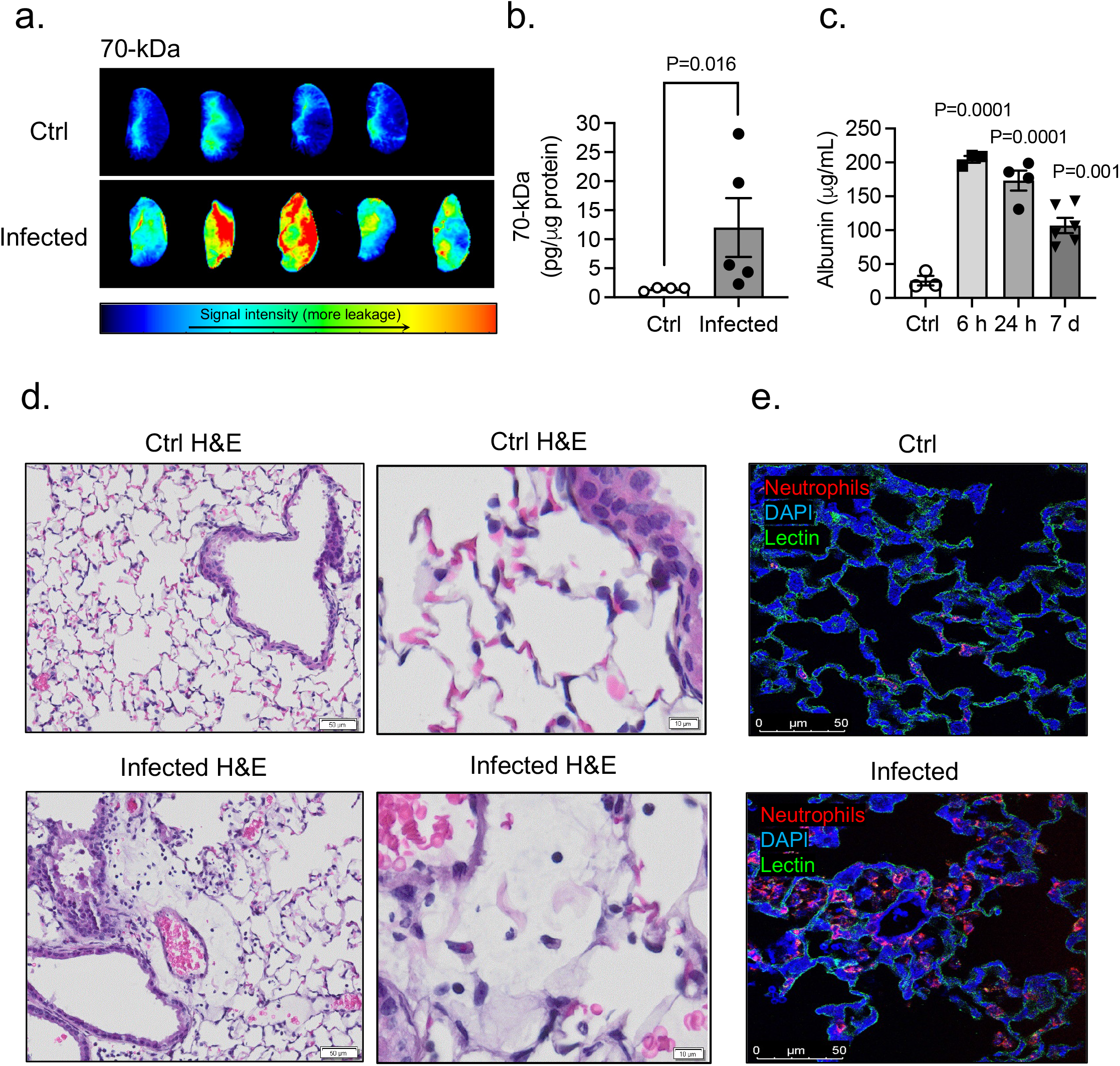
Effects of PA pneumonia on lung inflammation. (a) Representative NIR fluorescence images of the left lung lobe obtained from control and PA infected mice (OD_600_ 0.3) showing the distribution of 70-kDa tracer within the tissue. (b) Summary data showing lung permeability of 70-kDa tracer. Mann-Whitney test, P=0.016 vs. control (uninfected) group. (c) Plasma leakage indicated by increased levels albumin in bronchoalveolar lavage fluid (BALF). Kruskal-Wallis test, P=0.0001, P=0.001 vs. control (uninfected) group. (d) Histological analysis by conventional H&E staining of lung sections in control and PA infected mice (OD_600_ 0.3). (e) Serial lung sections obtained from control and PA-infected mice immunostained for neutrophils (red). Lectin (green) staining was used to visualized blood vessels and DAPI (blue) for nuclei. In all graphs, each point indicates data from an individual animal, bar graphs (columns and error bars) show mean ± S.E.M.

### BBB leakage following PA lung infection

Next, we studied whether PA-induced lung infection (OD_600_ 0.3) causes BBB permeability changes. BBB leakage was first evaluated 24 hours after infection. We used NIR imaging of whole brains after intravenous administration of 10-kDa and 70-kDa fluorescent tracers (Fig. 2a-d). Significant leakage of both tracers into the brain parenchyma was detected in the infected group. Importantly, paracellular permeability to 10-kDa dye was higher than the 70-kDa, suggesting a size-depending opening of the BBB (Fig. 2a,d). The time-course of BBB permeability changes (“BBB opening and recovery”) was evaluated with NaFl at 24 hours, 7 days, and 1-month after infection. Our results indicate a time-dependent increase in the degree of BBB hyperpermeability, which peaked at 24 hours after lung infection and decreased by ~80% by 1-month post-infection (Fig. 2e). The disruption of the BBB was associated with changes in mRNA expression of AJs and TJs in the brain at different time points post-infection (Fig. 2f). VE-cadherin mRNA levels decreased by day 7 and remained low 1-month after infection. Claudin-5 mRNA levels decreased at earlier time points (24-hours) after infection while occludin mRNA levels remained unchanged.

**Fig. 2.**
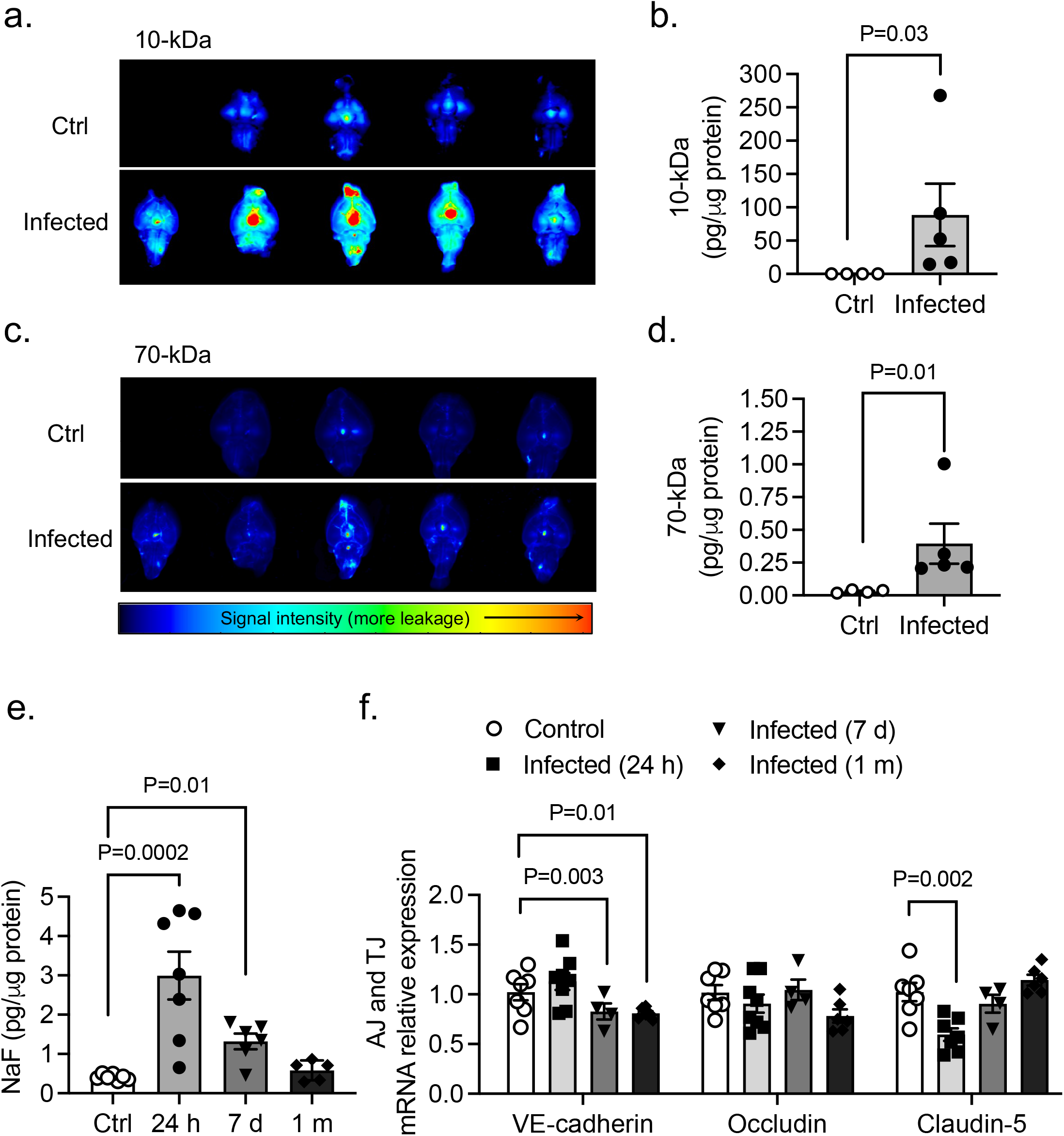
Effects of PA lung infection on BBB permeability and TJ expression. Representative NIR fluorescence images of whole brains obtained from control and PA infected mice (OD_600_ 0.3) showing the distribution of (a) 10-kDa and (c) 70-kDa tracers within the tissue. (b,d) Summary data showing BBB paracellular permeability of 10-kDa and 10-kDa fluorescence tracers. Mann-Whitney test, P<0.05 vs. control (uninfected) group. (e) Time course of BBB permeability changes to small size solutes as measured by NaFl uptake starting at 24 h post-infection to 1-month post-infection (OD_600_ 0.3). Kruskal-Wallis test, P=0.01 and P=0.0002 vs. control (uninfected) group. (f) Summary data showing mRNA expression levels of VE-cadherin, occludin and claudin-5 in brains harvested from control and PA infected mice at 24 hours, 7 days and 1 month after infection. Kruskal-Wallis test, P=0.01 and P=0.003 and P=0.002 vs. control (uninfected) group. In all graphs, each point indicates data from an individual animal, bar graphs (columns and error bars) show mean ± S.E.M.

### Lung infection by PA causes anxiety-like behaviors

Next, we assessed both groups of mice for locomotor and anxiety-like behavior in an open field 7 days after saline or PA inoculation (OD_600_ 0.3) (Fig. 3a). PA infection increased the total distance travelled in the open field (Fig. 3b) and decreased the amount of time animals remained immobile (Fig. 3c); the amount of time spent in the brightly-lit center of the open field was higher in infected animals compared to controls (Fig. 3d). The results of the open field test indicate PA-infected animals had increased locomotor activity and anxiety-like behavior preferring to stay in the center of the box compared with uninfected controls.

**Fig. 3.**
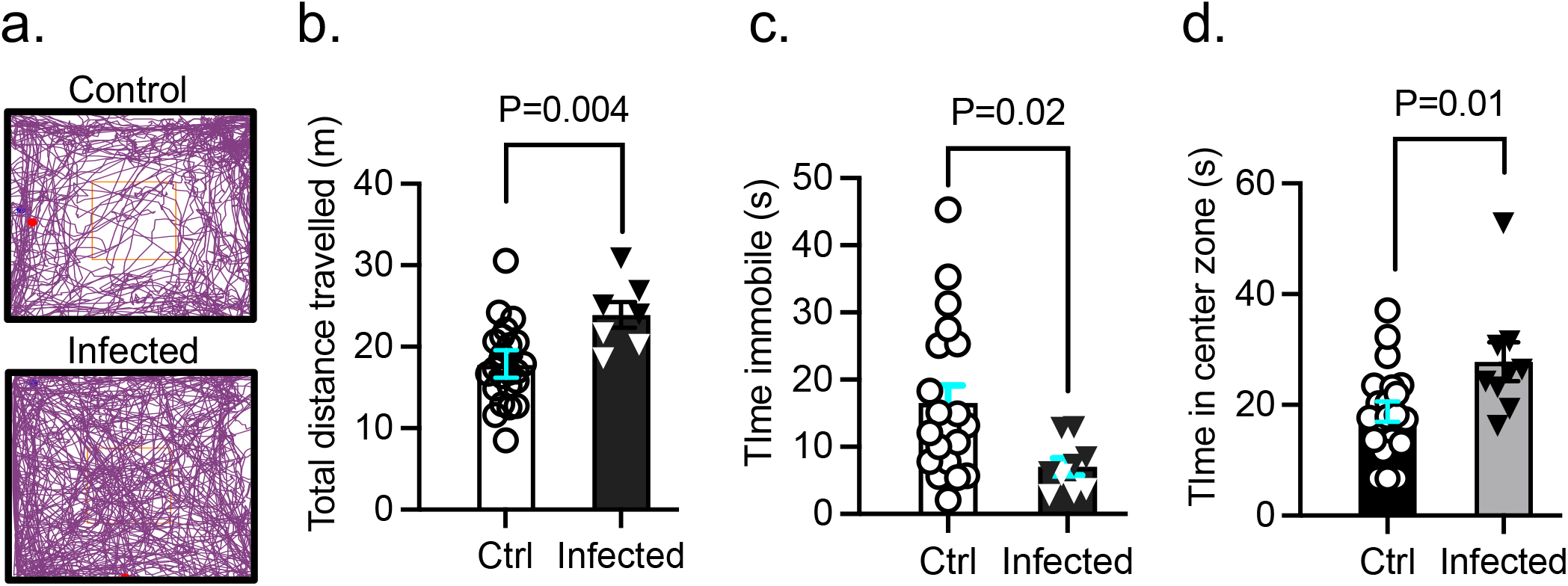
Effect of PA pneumonia on mouse behavior. (a) Representative tracks of control and PA-infected mice (OD_600_ 0.3) recorded by ANYMaze^®^ video system during the open field test. (b) Total distance traveled by control and PA-infected mice. (c) Time mice remained immobile in the open field. (d) Time spent in the center of the open field by controls and PA-infected mice. Mann-Whitney test, P=0.004, P=0.02, P=0.01 vs. control (uninfected) group. In all graphs, each point indicates data from an individual animal, bar graphs (columns and error bars) show mean ± S.E.M. In all graphs, each point indicates data from an individual animal, bar graphs (columns and error bars) show mean ± S.E.M.

### Tissue distribution of PA in vivo

We next characterized the spread of infection in mice intratracheally inoculated with PA (OD_600_ 0.3) 24 hours, 7 days, and 1-month after infection. We examined bacteria loads in the organs of infected mice by quantifying CFUs in lung and spleen homogenates and CSF (Fig. 4a). In the lungs, bacterial load decreased by ~4-fold 1-month after infection. PA load was also measured in the spleen, which was used as a peripheral organ to assess whether there is systemic dissemination of PA. In the spleen, PA was detected, although bacterial load was lower compared to lung at all time points post-infection (Fig. 4a). Next, we evaluated the presence of PA in the brain. We chose CSF for quantification of CFUs over homogenized brain tissue since ~90% of infected animals showed bacteremia 24 hours post-inoculation and brains could be contaminated with blood containing bacteria, which could provide misleading results. In parallel to the quantification of bacterial loads, we performed confocal microscopy to visualize GFP-labeled PA distribution within the lungs, spleen and brain. As shown in Figure 4b,c, 24 hours after infection PA was detected with abundance in the lungs, mainly around the airways and spleen. In the brain, we studied three putative sites of entry into the CNS: meningeal vessels, choroid plexus, brain parenchyma, hippocampus and isolated brain microvessels (Fig. 4d-g). We did not detect PA in any of those locations indicating the absence of PA infiltration into the CNS. Next, to confirm the lack of distribution of PA in live tissue, we performed multi-photon microscopy in freshly harvested lungs and brains 24 hours post-infection. The 3D reconstruction of z-stacks of brains and lungs confirmed results seen on fixed tissues. In the lungs, PA was localized throughout the lung, while no bacteria were detected in the brain parenchyma or inside brain capillaries (Suppl videos 1-4).

**Figure 4.**
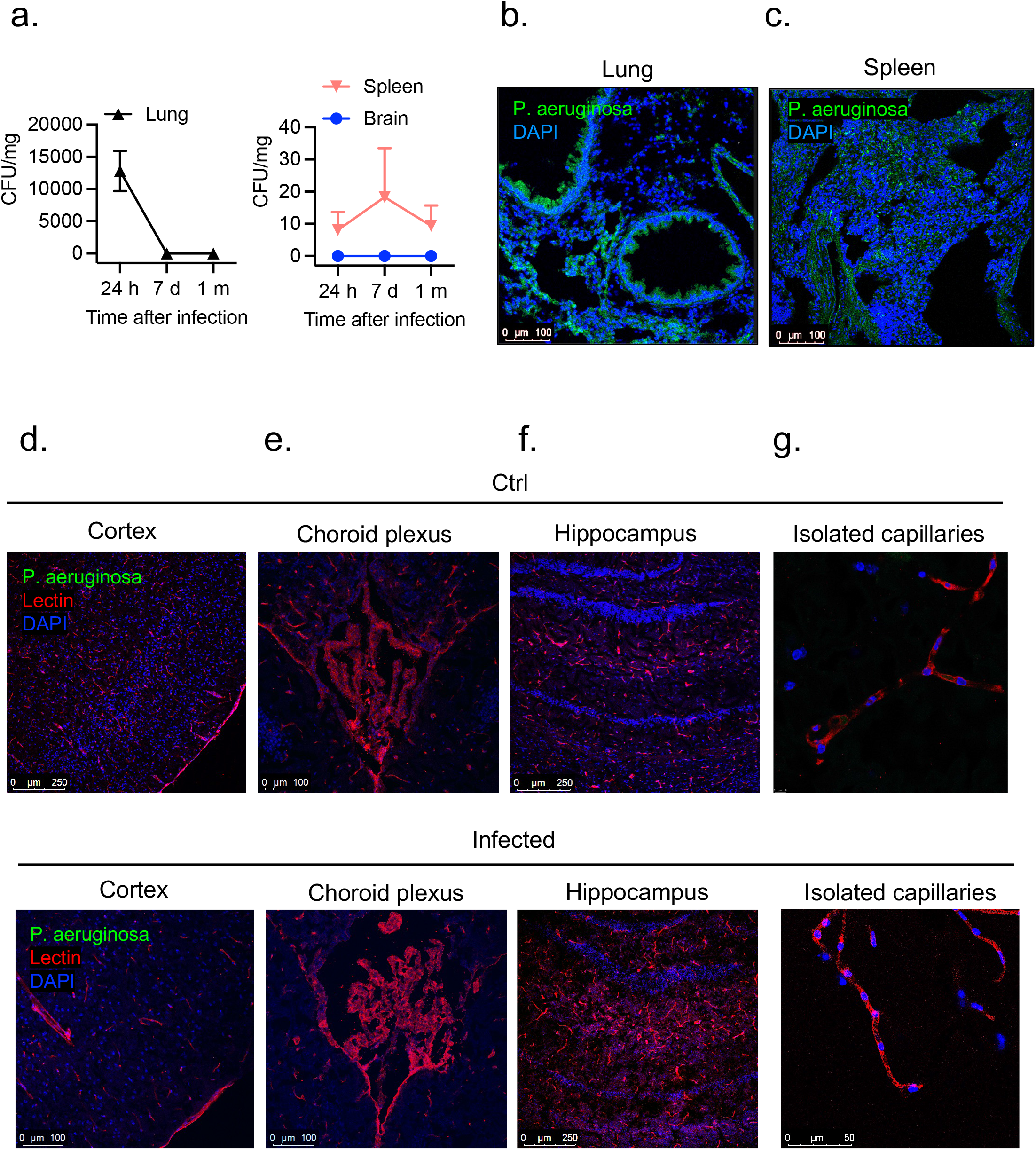
PA load in peripheral organs and brain regions. (a) Bacterial growth (CFUs per mg of tissue) in lung, spleen and in brain (CSF) obtained from infected animals (OD_600_ 0.3) and controls after intratracheal administration of PA. (b) Representative confocal micrograph showing (b) lung and (c) spleen sections obtained from GFP-labelled PA infected animals (OD_600_ 0.3) and controls. PA is labelled in green and nuclei (DAPI) in blue. (d-g) Representative confocal micrographs of (d) cortical sections, (e) choroid plexus (f) hippocampus and (g) isolated capillaries showing the lack of distribution of PA (GFP-labelled PA) in the brain of infected mice and controls. Lectin (red) was used to label the blood vessels and DAPI (blue) for the nuclei.

### Appearance of leukocytes in the CNS during PA pneumonia

We next investigated the potential systemic effects of PA-induced lung infection. PA inoculation (OD_600_ 0.3) into the lungs caused an increased level of total white blood cell count, along with significantly higher number of neutrophils and monocytes 24 hours post-infection (Fig. 5a,b). In addition, we studied whether PA pneumonia triggers an inflammatory response in the brain and evaluated whether there is infiltration of circulating leukocytes into the brain and changes in microglia number. In the CNS parenchyma, we found an abundant infiltration of leukocytes at 24 hours after infection. Subpopulations of leukocytes were discriminated according to the expression of CD11b and CD45: microglia (CD11b^+^ CD45^low^), myeloid leukocytes (CD11b^+^ CD45^hi^) and non-myeloid leukocytes (CD11b^-^ CD45^hi^) (for gating strategy see Suppl. Fig. 1). Analysis by flow cytometry confirmed elevated CD45^+^ CD11b^+^ cell numbers in the brain 24 hours post-infection and no change in microglia retrieved from brain tissue (Fig. 5c,d).

**Fig. 5.**
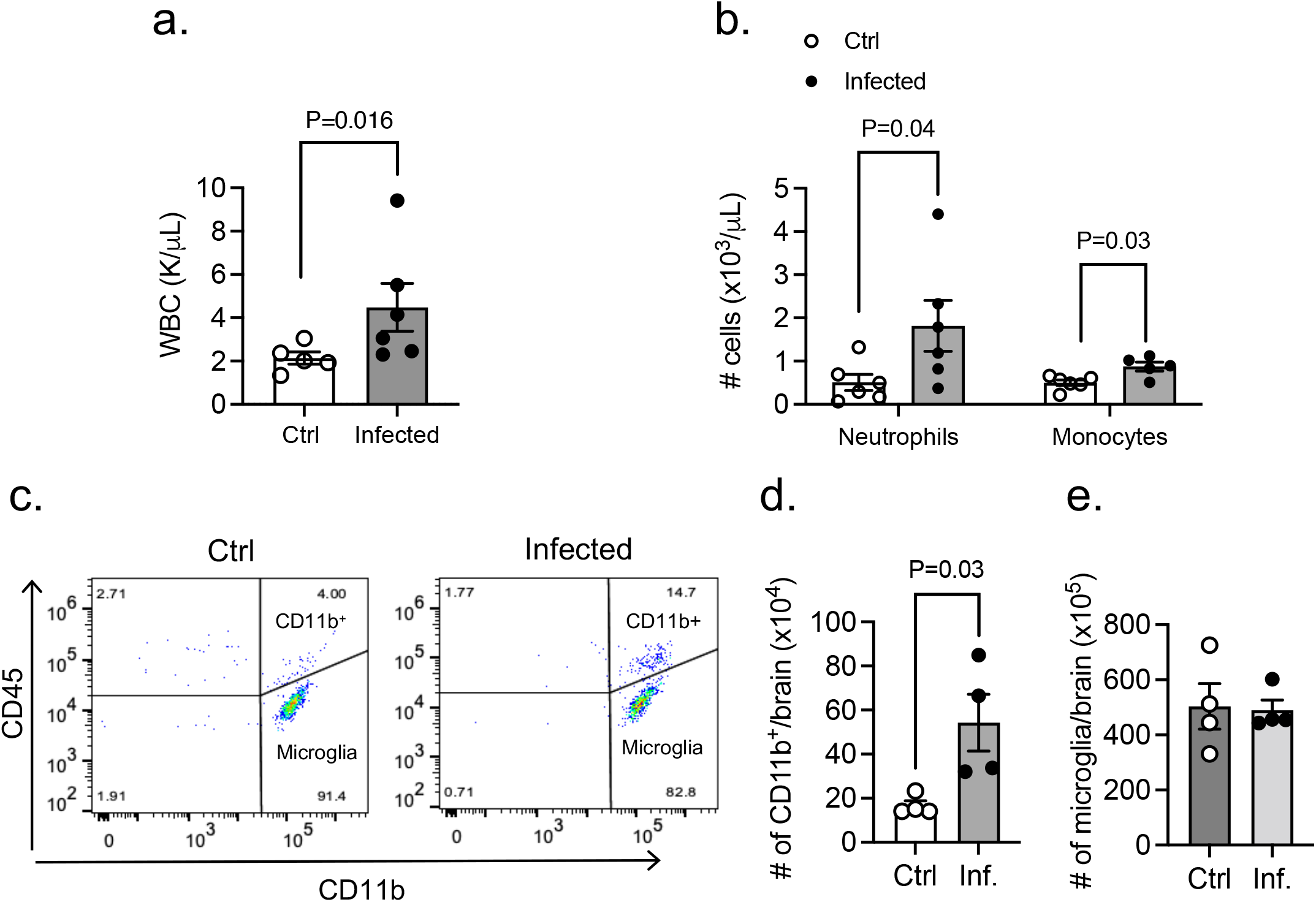
Appearance of leukocytes in CNS during PA-induced lung infection. (a) White blood counts in control and PA-infected animals (OD_600_ 0.3). Mann-Whitney test, P=0.016 vs. control (uninfected) group. (b) Neutrophil and monocyte counts in blood obtained from control and PA-infected mice (OD_600_ 0.3) measured by ProCyte One hematology analyzer. Mann-Whitney test, P=0.04, P=0.03 vs. control (uninfected) group. (c) Flow cytometry analysis of brain tissue homogenates labeled with anti-CD45 and anti-CD11b antibodies with gating on CD45^+^ cells. Quantitative analyses of (d) CD11b^+^ immune cell subpopulations and (e) microglia in control and PA infected mice. Mann-Whitney test, P=0.03 vs. control (uninfected) group. In all graphs, each point indicates data from an individual animal, bar graphs (columns and error bars) show mean ± S.E.M.

### PA pneumonia upregulates systemic cytokines and causes neuroinflammation

We next measured three representative systemic inflammation-derived proinflammatory cytokines (IL-6, IL-1β and TNF-α) following PA administration (OD_600_ 0.3). There was significant elevation of these proinflammatory cytokines in the plasma after lung infection peaking at early time points (6-24 hours) (Fig. 6a,c). We also examined the brain’s inflammatory state in vivo after PA lung infection. Notably, our results showed an upregulated transcriptional expression of cytokines (IL-1β, IL-6 and TNF-α), adhesion molecules (VCAM-1, ICAM-1) and chemokines (CXCL1 and CXCL2) after infection in both brain cortex (Fig. 6d-f) and hippocampus (Fig. 6g-i).

**Fig. 6.**
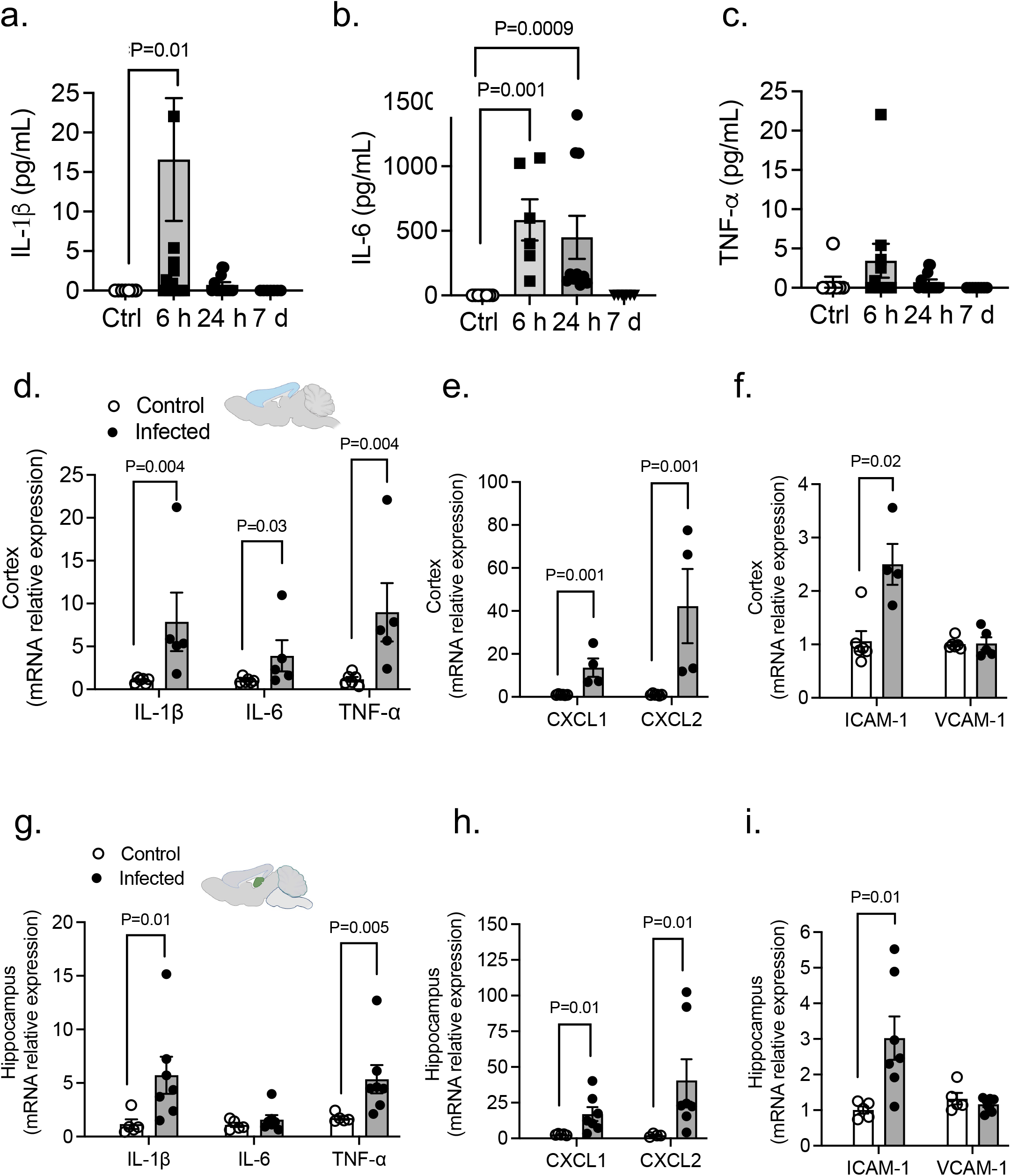
PA lung infection causes systemic and neuroinflammation. (a-c) PA-induced plasma levels of IL-6, IL-1β and TNF-α at different time points after infection (OD_600_ 0.3) compared to uninfected controls. Kruskal-Wallis test, P=0.01, P=0.001 and P=0.0009 vs. control (uninfected) group. (d-f) mRNA expression levels of (d) cytokines (IL-1β, IL-6 and TNF-α), (e) chemokines (CXCL1 and CXCL2) and (f) adhesion molecules (ICAM-1 and VCAM-1) in cortex from control and infected mice. Mann-Whitney test, P=0.004, P=0.03, P=0.001, P=0.001 vs. control (uninfected) group. (g-i) mRNA expression levels of (g) cytokines (IL-1β, IL-6 and TNF-α), (h) chemokines (CXCL1 and CXCL2) and (i) adhesion molecules (ICAM-1 and VCAM-1) in hippocampus from control and infected mice. Mann-Whitney test, P=0.01, P=0.005, P=0.01 vs. control (uninfected) group. In all graphs, each point indicates data from an individual animal bar graphs (columns and error bars) show mean ± S.E.M.

### Cytokine-induced brain endothelial barrier dysfunction

Cytokines are known to regulate not only immunity, but also barrier function (21–24). Once we determined that PA lung infection increases the level of circulating cytokines and there is no direct penetration of PA into the brain, we asked the question of whether circulating cytokines are responsible for the disruption of the brain endothelial barrier leading to BBB leakage. Brain microvascular endothelial cells were exposed for 24 hours to IL-1β (2, 20 and 200 ng/mL) and the combination of IL-1β and TNF-α (20 ng/mL). We elected those cytokines because they were highly expressed in the brain during PA pneumonia. Fig. 7a shows the impact of cytokine exposure on barrier tightness. For IL-1β, TER was significantly decreased by 50% (Fig. 7a,b), while the combination of cytokines caused a further decrease in TER (by 61%). In our previous work, brain microvascular endothelial cells showed a tight monolayer within 5 days, expressing ZO-1/claudin-5 and well-organized TJs, with a low permeability to small solutes (15). Next, we studied the localization of the TJ protein ZO-1 upon IL-1β treatment of brain endothelial cells (Fig. 7b). Endothelial cells under control conditions showed organized cell monolayers with clearly visible ZO-1 immunostaining. At the TER maximum (approximately 2.5 hours after PA challenge), TJ protein ZO-1 was still present outlining the outer of all cells under control conditions. After IL-1β treatment, ZO-1 immunostaining showed a diffused and discontinuous pattern with visible intercellular gaps, where ZO-1 immunostaining was not fully outlining the circumference of each cell, illustrating severe damage to the integrity of the endothelial cell layer.

**Fig. 7.**
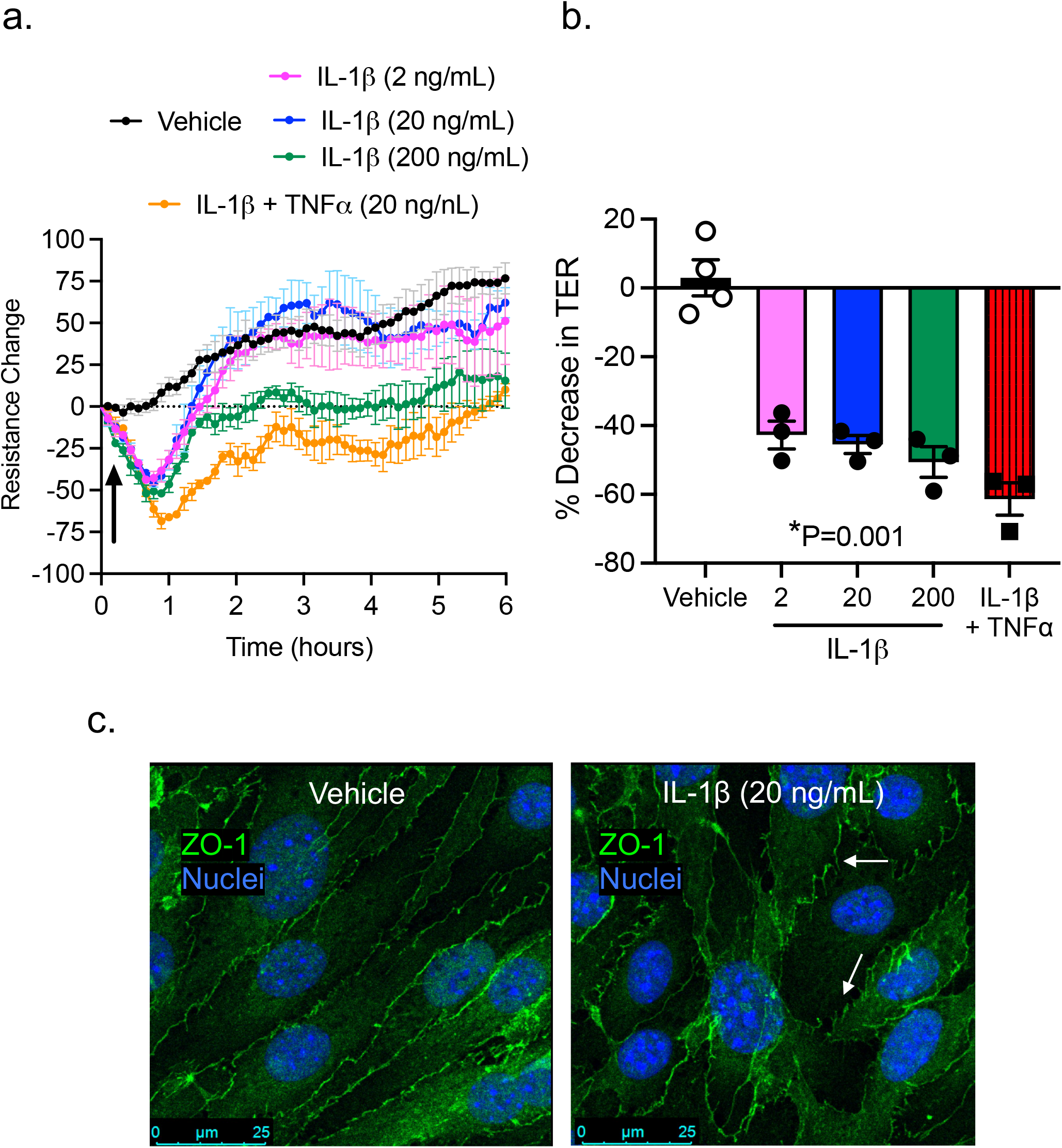
Effect of proinflammatory cytokines on brain microvascular endothelial barrier function. (a) TER measurements across confluent monolayers of mouse brain microvascular endothelial cells incubated with IL-1β (2-200 ng/mL) and IL-1β+TNF-α (20 ng/mL each). The data were represented as resistance change. Mean ±S.E.M., n=3. Arrow indicates when cytokines were added. (b) Summary data showing drop in TER values (peak). (c) Immunocytochemical analysis of ZO-1 (green) on brain microvascular endothelial cells under control conditions (vehicle) or IL-1β treatment in vitro. Nuclei was stained with DAPI. White arrows indicate TJ disorganization.

## Discussion

This study shows that PA pneumonia causes BBB leakage, neuroinflammation, and behavior alteration (locomotor hyperactivity). The brain dysfunction occurred in the absence of direct infection of the cerebral vasculature or brain parenchyma; thus, we propose that the underlying mechanism involves an inflammatory response in the lung-brain axis linked by the circulation, characterized by an increased expression of cytokines/chemokines and higher number of leukocytes in the blood and CNS. This is further supported by our data showing that cytokines exerted direct effects on cerebral microvascular endothelium to disrupt TJs, reduce cell-cell adhesive barrier resistance, and increase BBB paracellular permeability.

PA is an opportunistic pathogen that can cause severe infectious disease in critically ill or immunocompromised individuals (25). PA is responsible for a high percentage of hospital-acquired pneumonia in patients on medical devices, such as ventilators or catheters, as well as patients undergoing major surgeries (14, 26, 27). Compared to the lungs and other organs, brain infection by PA is less common and limited to patients with head trauma or undergoing neurosurgical procedures, such as intracranial catheterization where PA gains direct access to the CNS (28, 29). In this study, we used a clinically relevant mouse model of pneumonia via intratracheal delivery of PA. The animals received a moderate-to-high level of bacteria, which resulted in ~50% mortality at 7 days post-infection. This is in agreement with a previous study showing that administration of PA into murine lungs led to severe sepsis and death (30). In our study, infected mice who survived for at least 24 hours demonstrated typical symptoms of acute pneumonia, including low body temperature, high white blood cell and neutrophil counts, acidosis, and proteinemia. Histopathological observations indicated that intratracheal inoculation of PA triggered an inflammatory response in the lungs characterized by alveolar-capillary membrane thickening and neutrophil infiltration. These changes coincided with pulmonary microvascular thrombosis, damaged endothelial lining, and capillary leakage. Infected animals showed high bacteria load in the lungs and blood stream. Together, these data support the validity of the PA-induced pneumonia model.

Interestingly, while PA positive cultures were obtained in the spleen, an organ remote from the lung, CSF cultures were negative. The absence of bacteria in the CNS is further supported by immunofluorescence and multiphoton microscopic evidence confirming that in our murine model of pneumonia, PA did not gain direct access to the brain. This finding is consistent with many other studies showing that Pseudomonas meningitis is often post-neurosurgical, rather than through direct penetration of PA into the CNS (26, 31, 32). On the other hand, lung infection triggers a rapid and orchestrated activation of inflammatory cells and pathways leading to the production of an array of inflammatory mediators. These mediators can target tissues remote from the initial site of infection and cause multiple organ dysfunction. Within this context, the pathophysiological importance of the inflammatory response in the lung-brain axis to bacterial infection has been well recognized. For example, recent studies have identified lung amyloids, tau proteins and endothelial glycocalyx fragments as bioproducts of acute lung infection can reach the CNS and affecting learning and memory (33–36). However, the molecular mechanisms underlying lung-brain interactions remain poorly understood. In particular, whether and how bacteria invasion into the lung affects the BBB structure and function have not been well studied. The present study provides novel evidence for the effects of PA lung infection on BBB permeability.

The maintenance of brain homeostasis is largely controlled by the BBB, a specialized neurovascular structure based on tightly sealed endothelial cells connected via cell-cell junctions (e.g., occludin, claudin, cadherin) surrounded by pericytes and astrocytic end-feet (37). The BBB exhibits a very low permeability and is highly restrictive to the passage of cells, solutes, and pathogens, thus preventing neurotoxicity (38). During pathological states such as injury or systemic inflammation, BBB breakdown leads to neuronal damage by allowing a plethora of circulating agents to enter the brain. Our data suggest that the BBB function is compromised during lung bacterial infection. In vivo, infected animals showed BBB leakage to tracers of different sizes, especially small molecules (NaF and 10-kDa), which indicates an increased paracellular permeability via compromised cell-cell junctions. Indeed, further molecular analyses showed a decreased expression of claudin-5 (TJ) and VE-cadherin (AJ). We also observed a differential response of these junction molecules at different times. For example, while claudin-5 levels started to decline early (within 24 hours) post-infection, VE-cadherin decreased most significantly during the late stage (days) after infection. In addition, the expression level of occludin remained unaltered throughout the time course. There has been controversy regarding the role of occludin in determining TJ strength and BBB permeability property; while one study showed that occludin knockout mice present intact BBB (39), others have reported that changes in occludin expression induced by proinflammatory cytokines lead to changes in paracellular permeability (40, 41).

As the endothelial barrier is in contact with circulating mediators whose levels vary under different pathophysiological conditions (42), we sought to characterize the direct effects of proinflammatory cytokines on BBB junction structure and function. Our model showed high levels of proinflammatory cytokines in the plasma of infected animals, supporting that systemic inflammation is part of disease progression following PA lung infection. In particular, the plasma levels of TNF-α, IL-1β and IL-6 were significantly elevated. Importantly, we also detected an increased expression of multiple cytokines and chemokines in the brain, with IL-1β and TNF-α as the most significantly elevated in both cortex and hippocampus regions. Therefore, we selected these cytokines for testing their effects on brain microvascular endothelial junctions in vitro. Our barrier resistance (ECIS) data showed that IL-1β induced a dose-dependent reduction in TER, indicator of cell-cell junction function, and this effect was augmented during combined treatment with IL-1β and TNF-α. Structurally, compared to control brain endothelial monolayers showing clearly defined TJs with continuous lining of ZO-1 at cell-cell contacts, IL-1β-treated endothelial cells showed discontinuous junction lining with ZO-1 stained in a finger-like pattern and intercellular gaps. These results reinforce the cytokines’ ability to disrupt the BBB integrity.

Indeed, previous studies have demonstrated a correlation between in vivo administration of pro-inflammatory cytokines and BBB leakage (39, 43). The ability of cytokines to open the paracellular pathway of endothelial barrier has been attributed to endothelial cytoskeleton reorganization (24) and/or downregulation of junction protein expression (22, 40, 44–50).

The microvascular leakage and BBB hyperpermeability observed in PA-infected animals were associated with neuroinflammation, evidenced by immune cell recruitment and high expression of proinflammatory cytokines, chemokines and adhesion molecules in the brain of mice with pneumonia. It is known that lung infection or injury is associated with various neurological complications, from altered memory to cognitive decline (51–54). In our model, even in the absence of direct penetration of PA into the brain, infected mice developed anxiety-like behaviors, which might result from cerebral microvascular dysfunction, BBB leakage, and systemic/local production of cytotoxic cytokines like IL-1β (55–57) and TNF-α (47, 58) as previously documented.

## Conclusions

The present study demonstrates that bacterial infection in the lungs causes neuroinflammation, cerebral microvascular leakage and increased BBB paracellular permeability; the underlying mechanisms involve cytokine-induced brain endothelial cell-cell adhesive barrier dysfunction and TJ downregulation. These data provide novel evidence supporting the mechanistic linkage between the lung and brain as part of an orchestrated host immune response to bacterial infection. In addition, our study is unique because we have developed and validated a mouse model of pneumonia that may prove useful in studying the brain sequelae of infectious diseases originated from non-CNS organs.

## Supporting information

Sppl Figs

## List of abbreviations

AJs: Adherens junctions
BBB: Blood-brain barrier
CFUs: Colony forming units
CNS: Central Nervous System
ELISA: Enzyme-linked immunosorbent assay
GFP: Green fluorescent protein
H&E: Hematoxylin and Eosin
IL: Interleukin
NaFl: Sodium fluorescein
OD: Optical density
PA: Pseudomonas aeruginosa
PBS: Phosphate buffer saline
PFA: Paraformaldehyde
TER: Transendothelial electrical resistance
TJs: Tight junctions
TNF: Tumor necrosis factor

## Declarations

## Competing interests

The authors declare that they have no competing interests

## Funding

This work was supported by the National Institutes of Health grants HL150732 and GM142110 (to SYY).

## Author’s contributions

NV performed experiments, analyzed data and wrote the manuscript. YM and SS performed flow cytometry experiments. SAG performed experiments, PA cultures and helped with data analysis. AJA and KN helped with neurological testing experiments. XY helped with tissue/blood collection. SYY conceptualize the study, edited and revised the manuscript and provided support through all levels of development. All authors approved the final version of the manuscript.

## Acknowledgements

No applicable

## Additional files.

**Suppl. Fig. 1.** (a) Representative flow cytometry plots illustrate the gating strategy used for this experimental series: single cell population (top left) was gated based on forward-scatter characteristics, Zombie viability dye was used to exclude dead cells (high fluorescence), and CD45 and CD11b staining were used to identify myeloid leukocytes (CD11b+ CD45^hi^) and microglia (CD11b+ CD45^low^) subpopulation of cells.

**Suppl. Fig. 2. Effect of PA on mouse survival, body temperature and biochemistry parameters.** (a) Survival rates of mice treated with PA at OD_600_ 0.3 and 0.6. (b) Body temperature obtained by measuring the rectal temperature after PA infection (indicated by arrow). (c) Blood pH values obtained from control and PA-infected mice at 4 hours and 24 hours after infection measured with iSTAT. (d) Total protein (albumin and globulin) concentration in blood measured in controls and 24-hours after PA infection.

## Notes

### Competing Interest Statement

The authors have declared no competing interest.

