## Supplementary figures and images for "Lung infection by *P. aeruginosa* induces neuroinflammation and blood-brain barrier dysfunction in mice"

### Sppl Figs

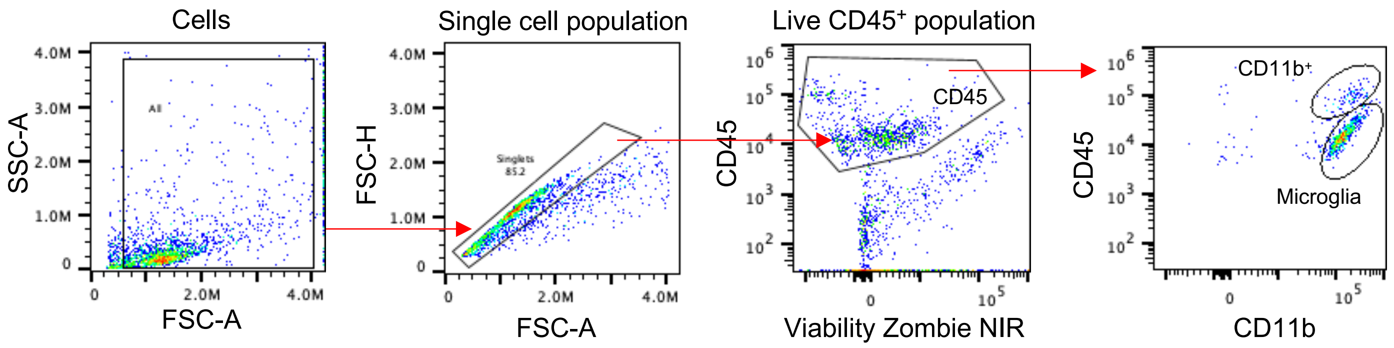

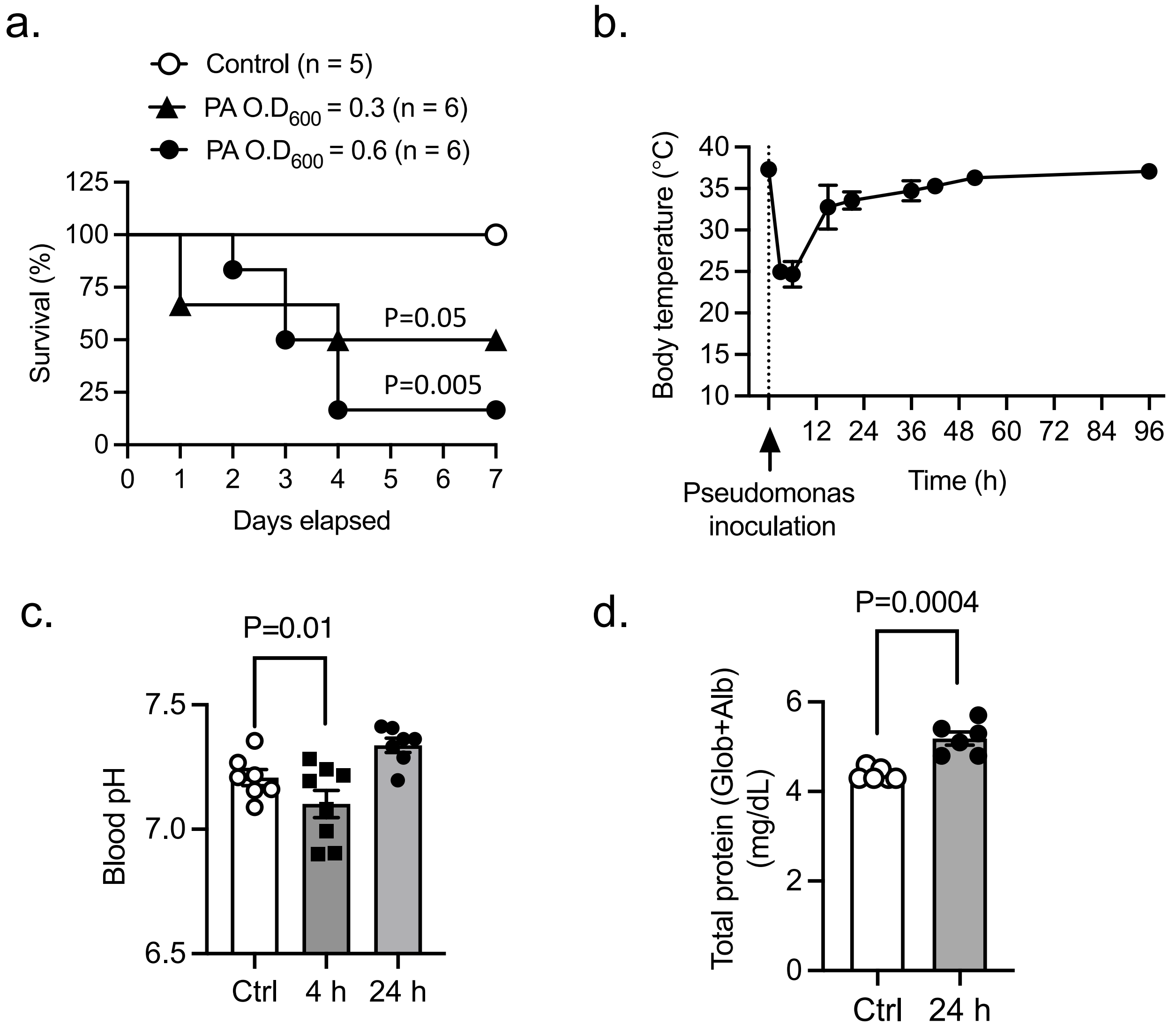
